## Supplemental figure for "Development of a new 3D tracking system for multiple marmosets under free-moving conditions"

### Slide 1
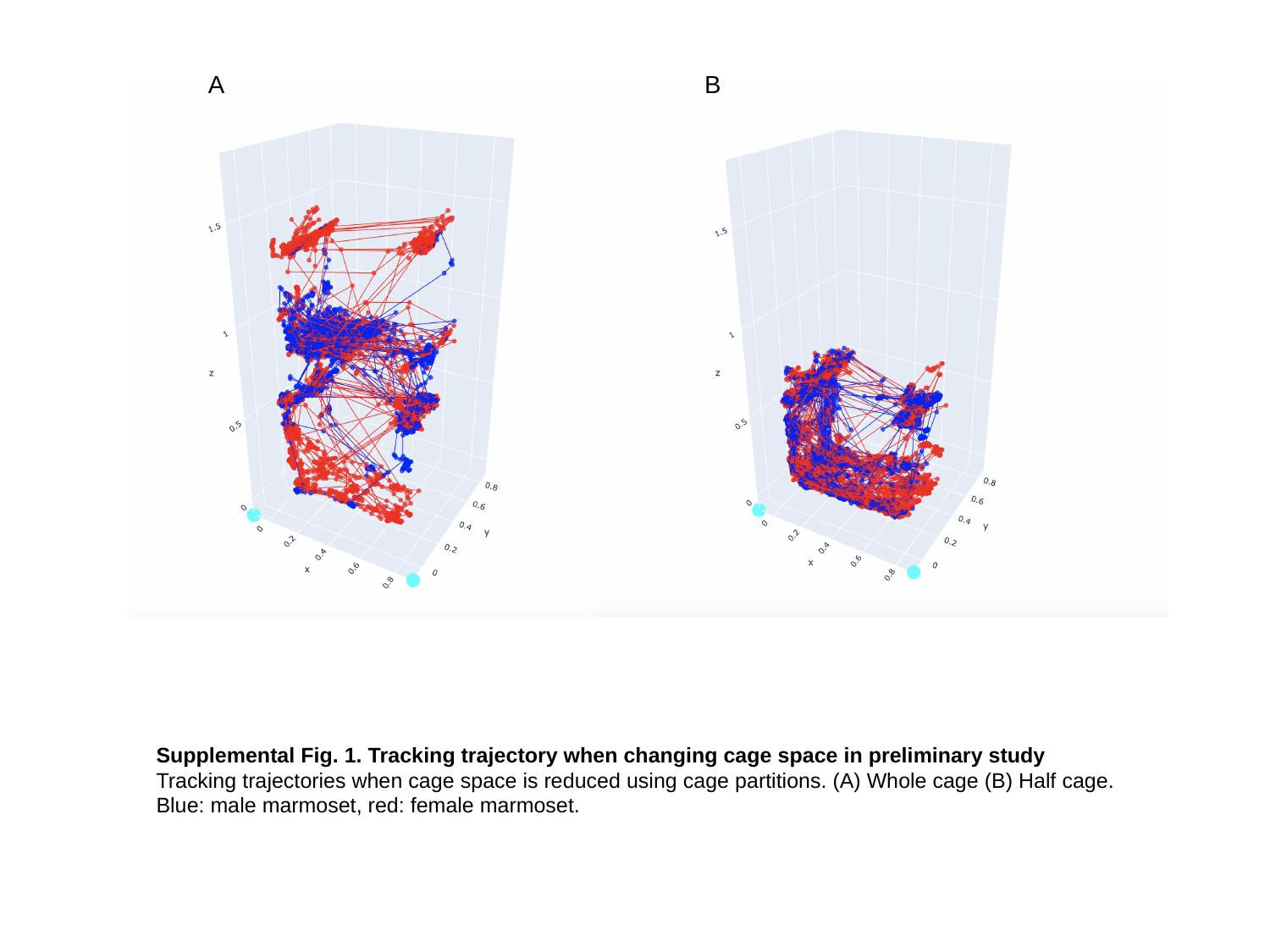

A
B
Supplemental Fig. 1. Tracking trajectory when changing cage space in preliminary study
Tracking trajectories when cage space is reduced using cage partitions. (A) Whole cage (B) Half cage. Blue: male marmoset, red: female marmoset.

### Slide 2
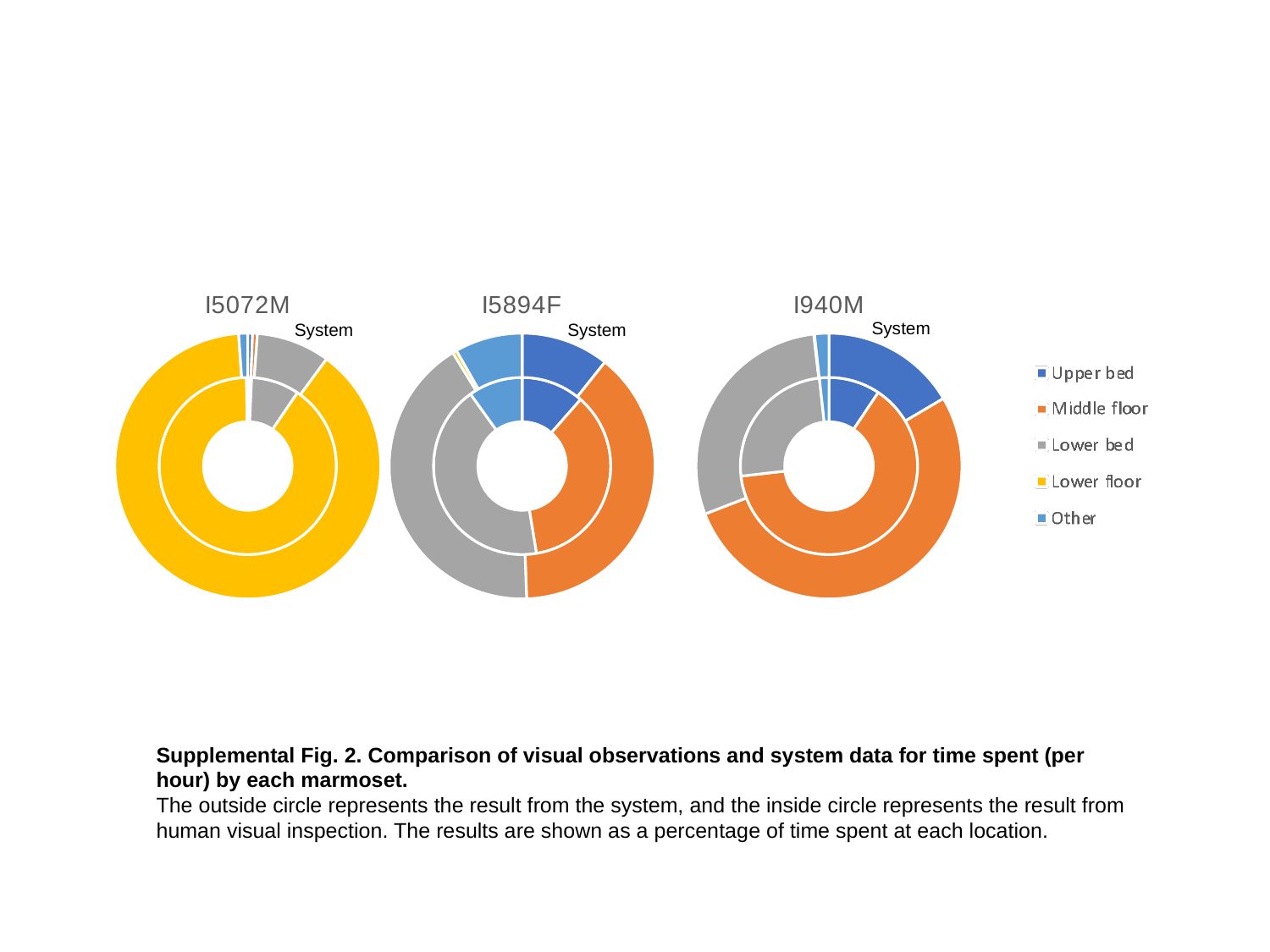

#### Chart: I5072M
| Category | | |
|---|---|---|
| Upper bed | 0.0036666666666683055 | 0.0058 |
| Middle floor | 0.0033333333333327884 | 0.00585 |
| Lower bed | 0.08766666666667185 | 0.0894 |
| Lower floor | 0.9030000000000005 | 0.8879 |
| Other | 0.002333333333326637 | 0.011050000000000004 |
#### Chart: I5894F
| Category | | |
|---|---|---|
| Upper bed | 0.11466666666666966 | 0.10708 |
| Middle floor | 0.35933333333334044 | 0.3875 |
| Lower bed | 0.4269999999999996 | 0.41813 |
| Lower floor | 0.0 | 0.005 |
| Other | 0.09899999999999032 | 0.08228999999999997 |
#### Chart: I940M
| Category | | |
|---|---|---|
| Upper bed | 0.0940000000000019 | 0.16513 |
| Middle floor | 0.6376666666666555 | 0.526037 |
| Lower bed | 0.25100000000000405 | 0.2905 |
| Lower floor | 0.0 | 0.0009 |
| Other | 0.01733333333333853 | 0.017433000000000032 |
System
System
System
Human
Human
Human
Supplemental Fig. 2. Comparison of visual observations and system data for time spent (per hour) by each marmoset.
The outside circle represents the result from the system, and the inside circle represents the result from human visual inspection. The results are shown as a percentage of time spent at each location.

### Slide 3
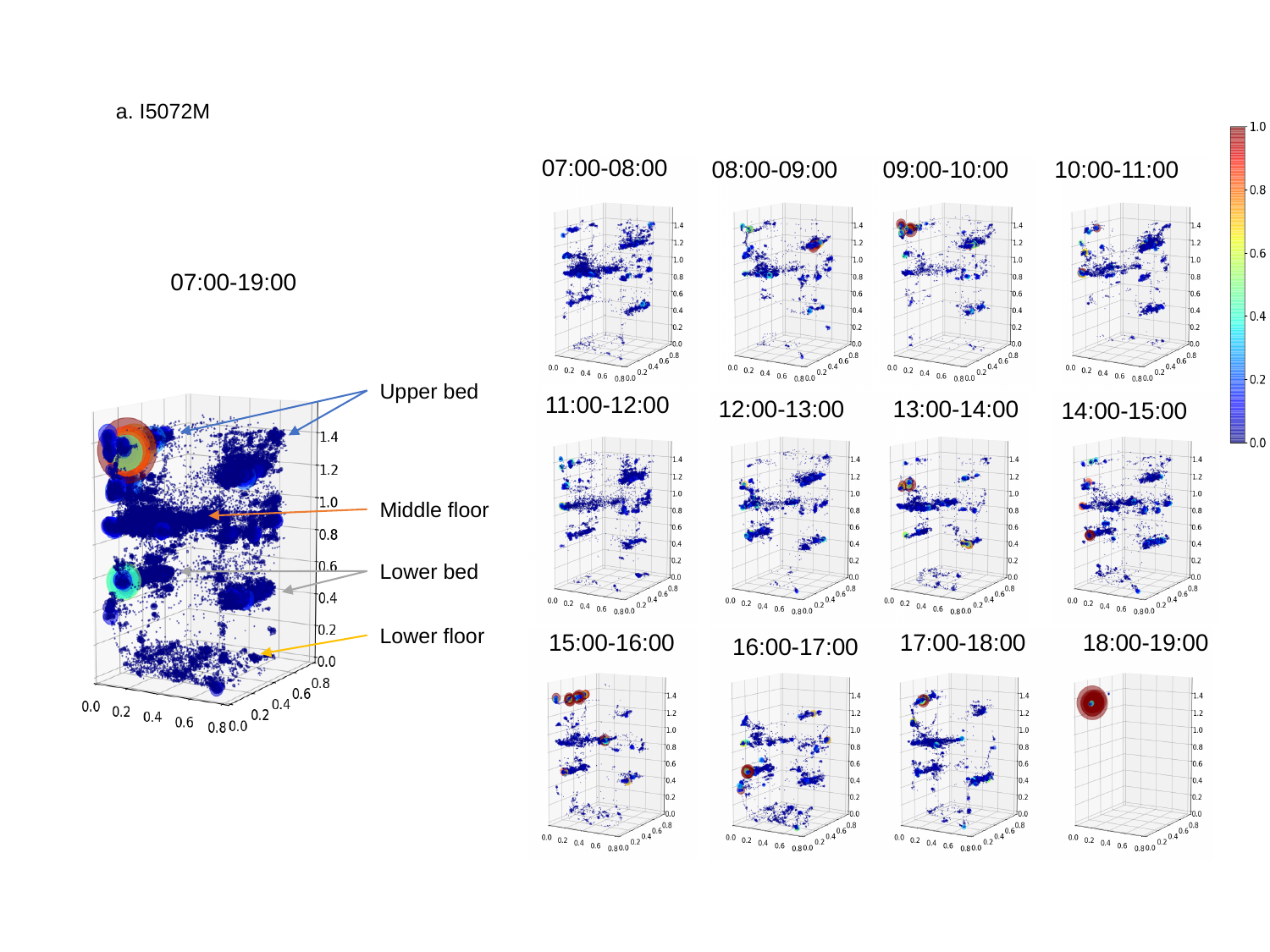

07:00-08:00
08:00-09:00
10:00-11:00
09:00-10:00
07:00-19:00
Upper bed
11:00-12:00
12:00-13:00
13:00-14:00
14:00-15:00
Middle floor
Lower bed
Lower floor
15:00-16:00
17:00-18:00
18:00-19:00
16:00-17:00
a. I5072M

### Slide 4
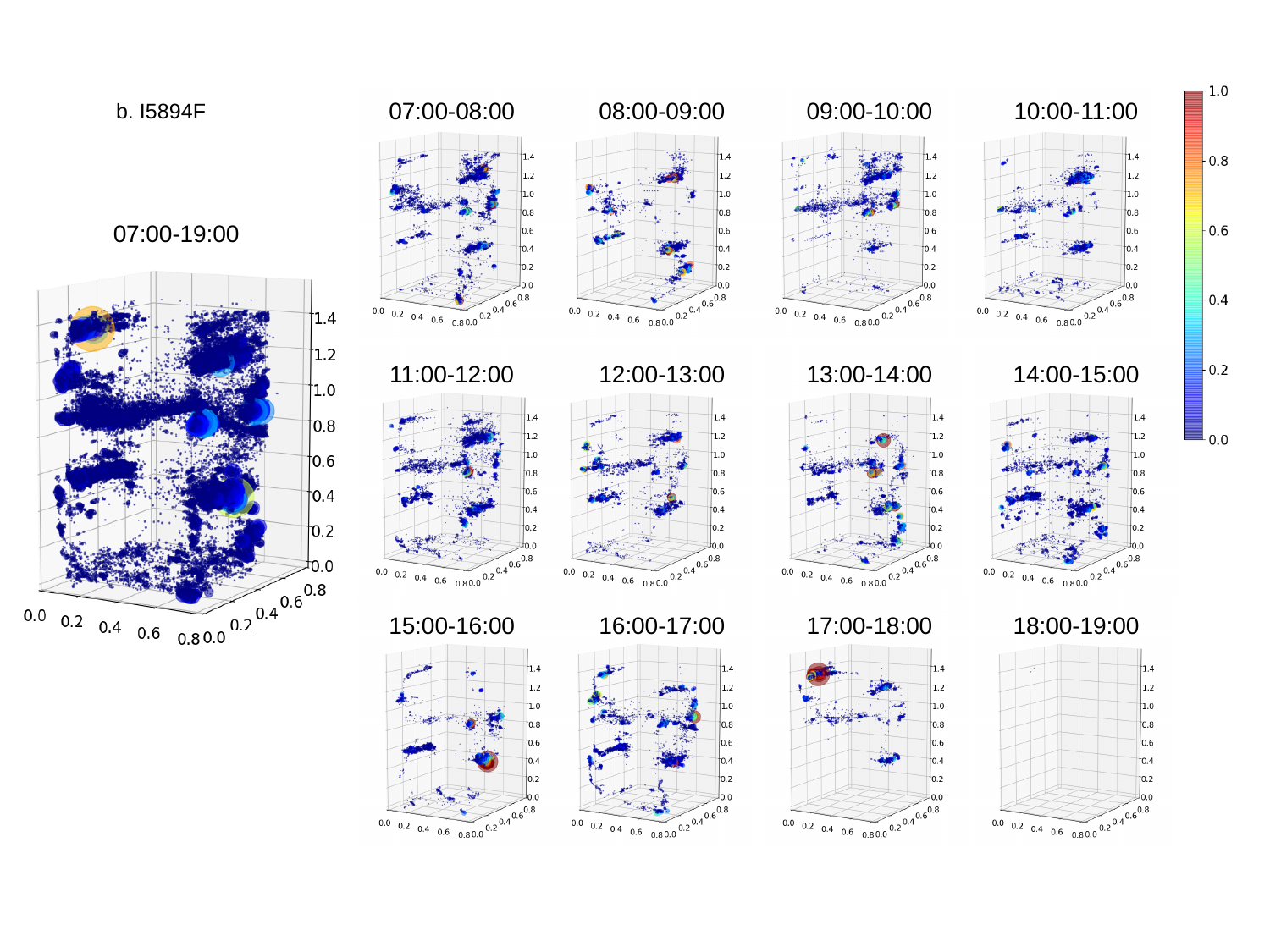

07:00-08:00
08:00-09:00
09:00-10:00
10:00-11:00
11:00-12:00
12:00-13:00
13:00-14:00
14:00-15:00
15:00-16:00
16:00-17:00
17:00-18:00
18:00-19:00
b. I5894F
07:00-19:00

### Slide 5
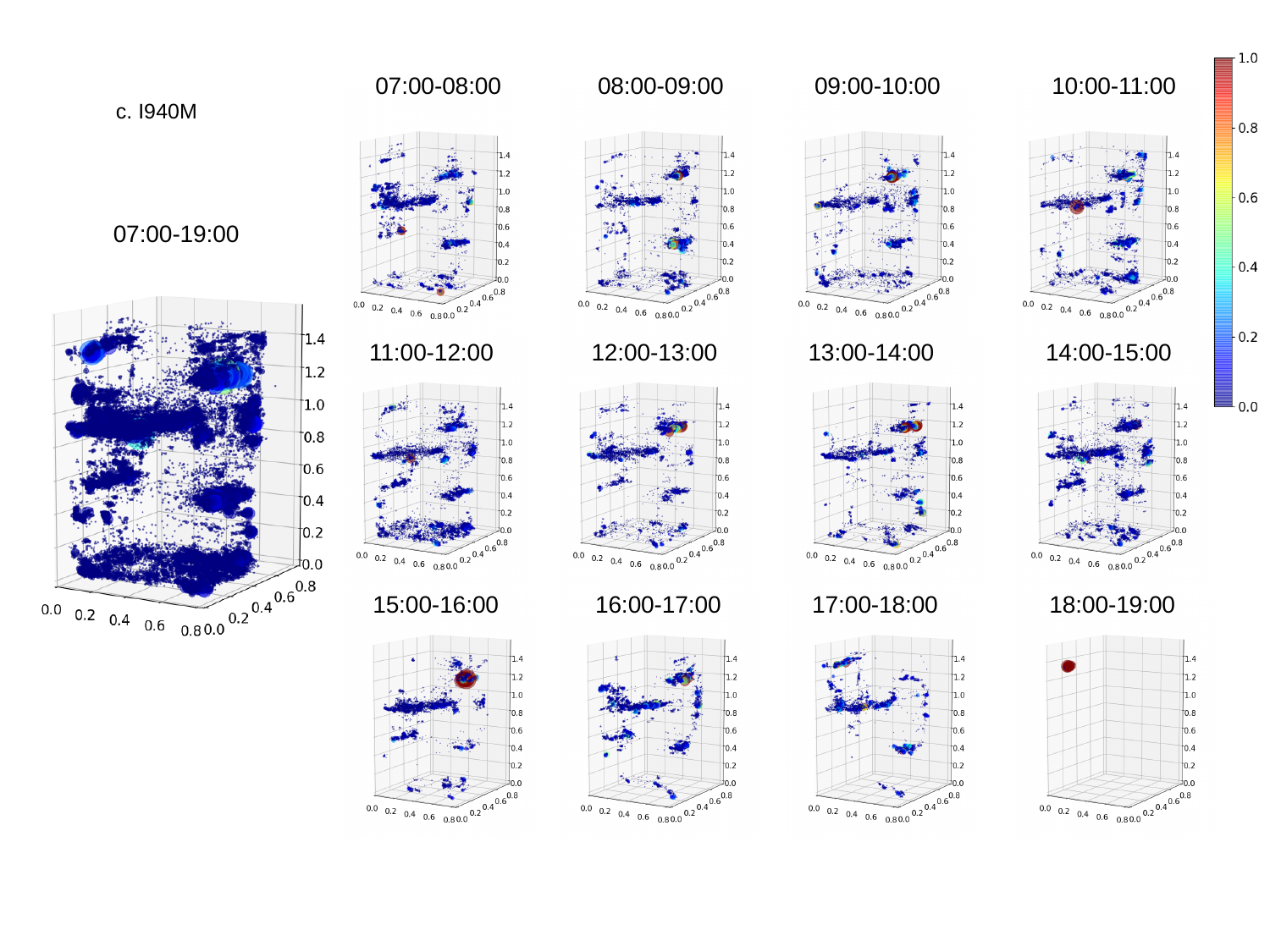

07:00-08:00
08:00-09:00
09:00-10:00
10:00-11:00
c. I940M
07:00-19:00
11:00-12:00
12:00-13:00
13:00-14:00
14:00-15:00
15:00-16:00
16:00-17:00
17:00-18:00
18:00-19:00

### Slide 6
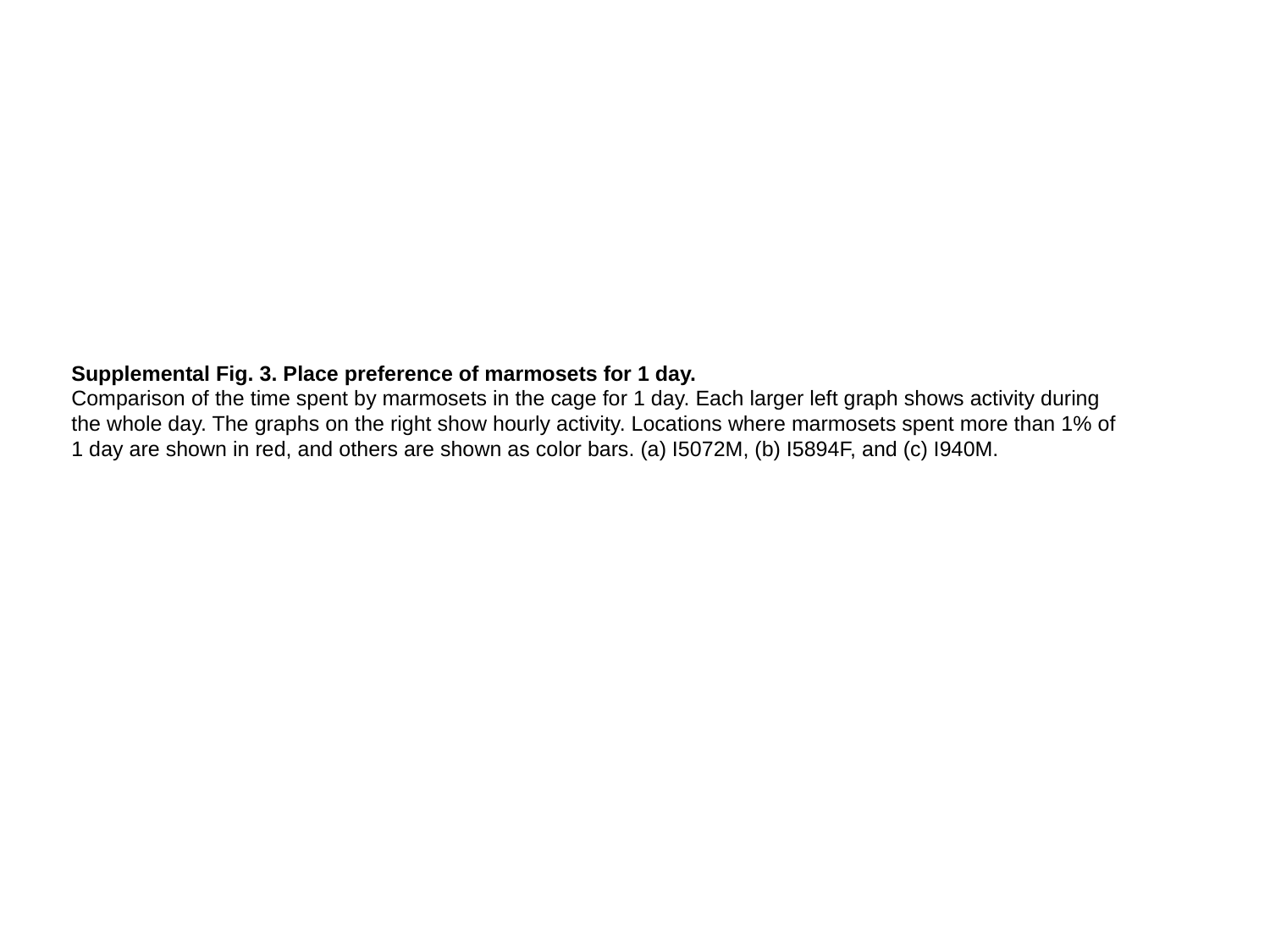

Supplemental Fig. 3. Place preference of marmosets for 1 day.
Comparison of the time spent by marmosets in the cage for 1 day. Each larger left graph shows activity during the whole day. The graphs on the right show hourly activity. Locations where marmosets spent more than 1% of 1 day are shown in red, and others are shown as color bars. (a) I5072M, (b) I5894F, and (c) I940M.

### Slide 7
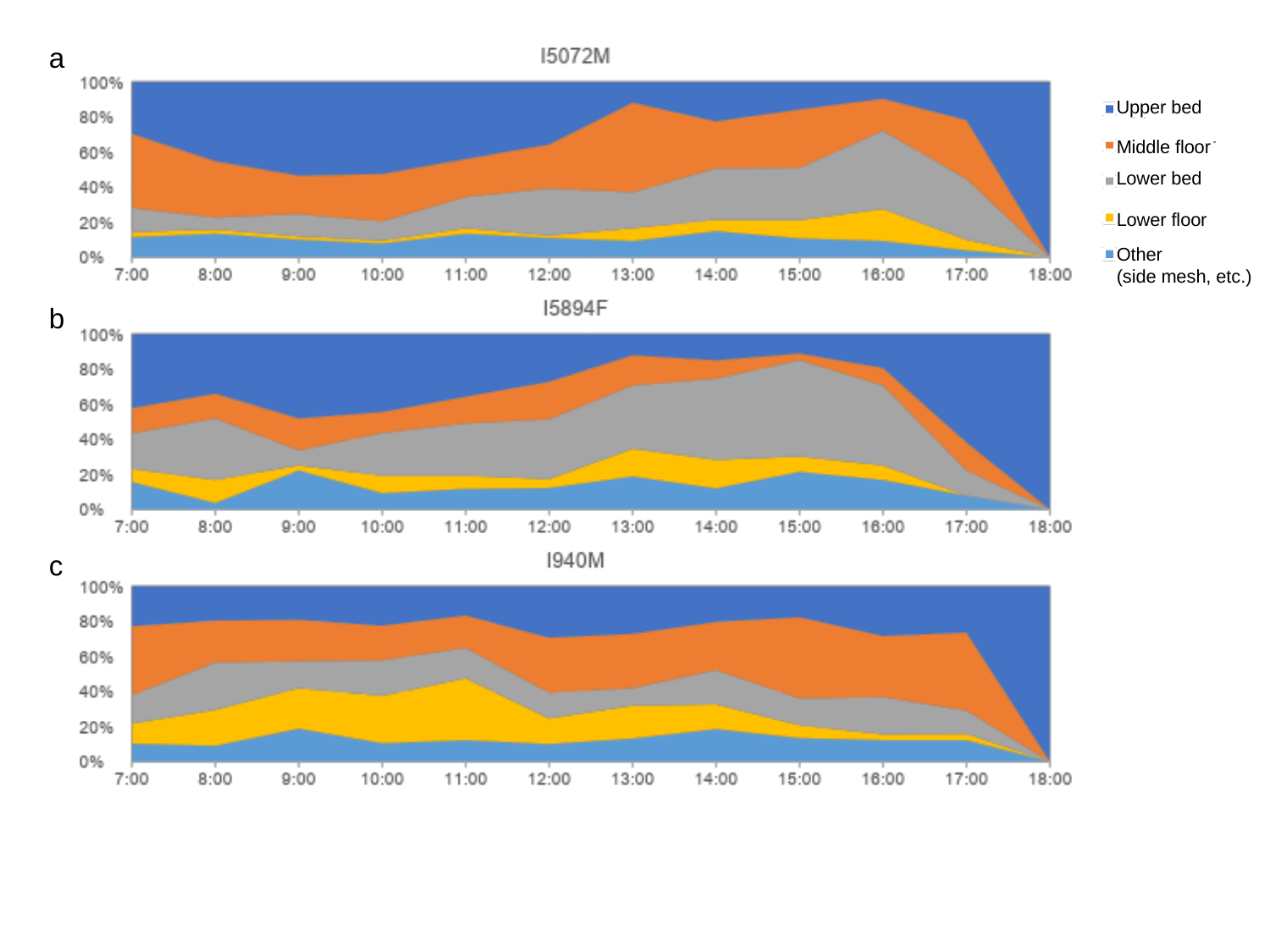

a
Upper bed
Middle floor
Lower bed
Lower floor
Other
(side mesh, etc.)
b
c

### Slide 8
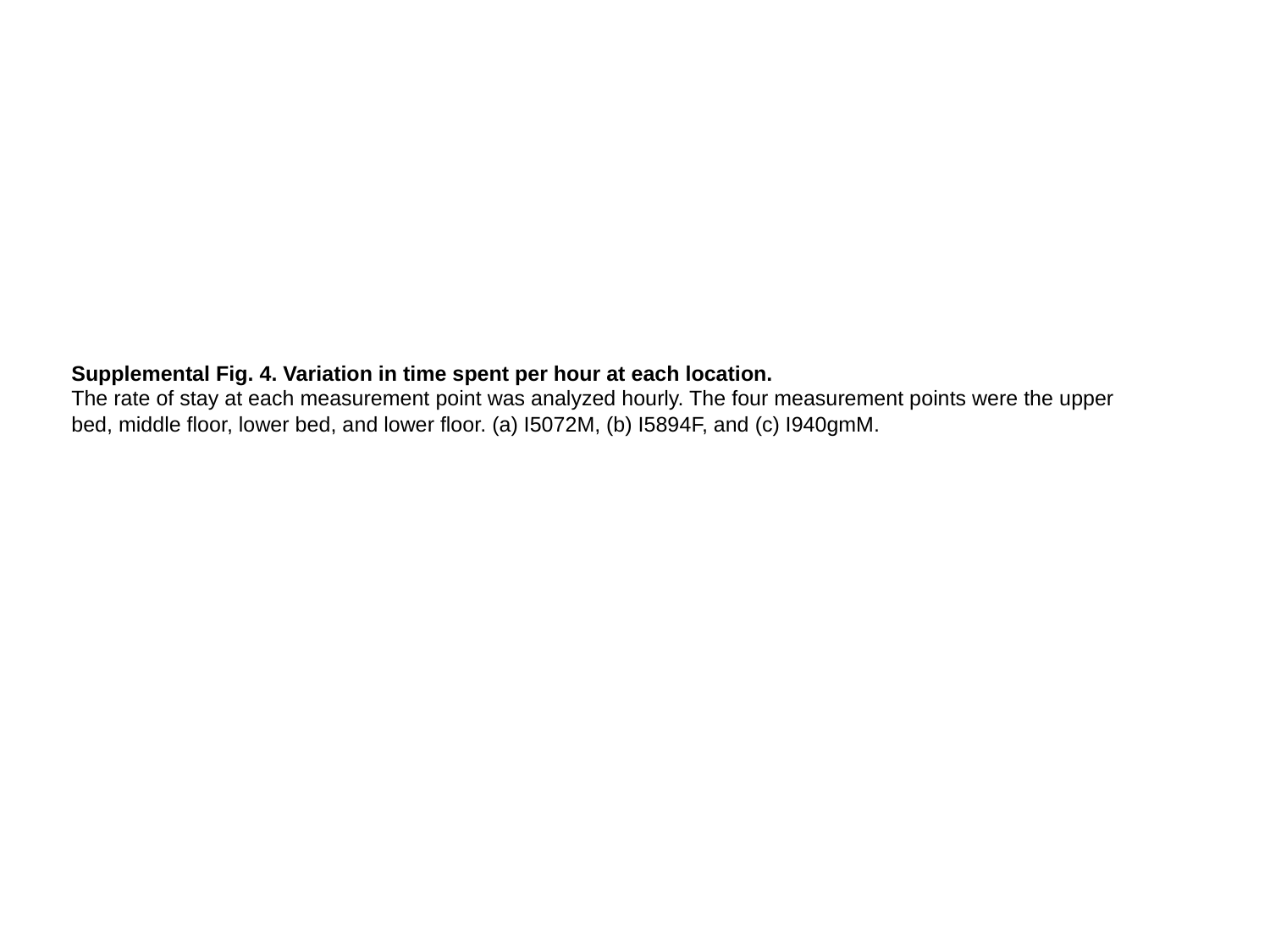

Supplemental Fig. 4. Variation in time spent per hour at each location.
The rate of stay at each measurement point was analyzed hourly. The four measurement points were the upper bed, middle floor, lower bed, and lower floor. (a) I5072M, (b) I5894F, and (c) I940gmM.

### Slide 9
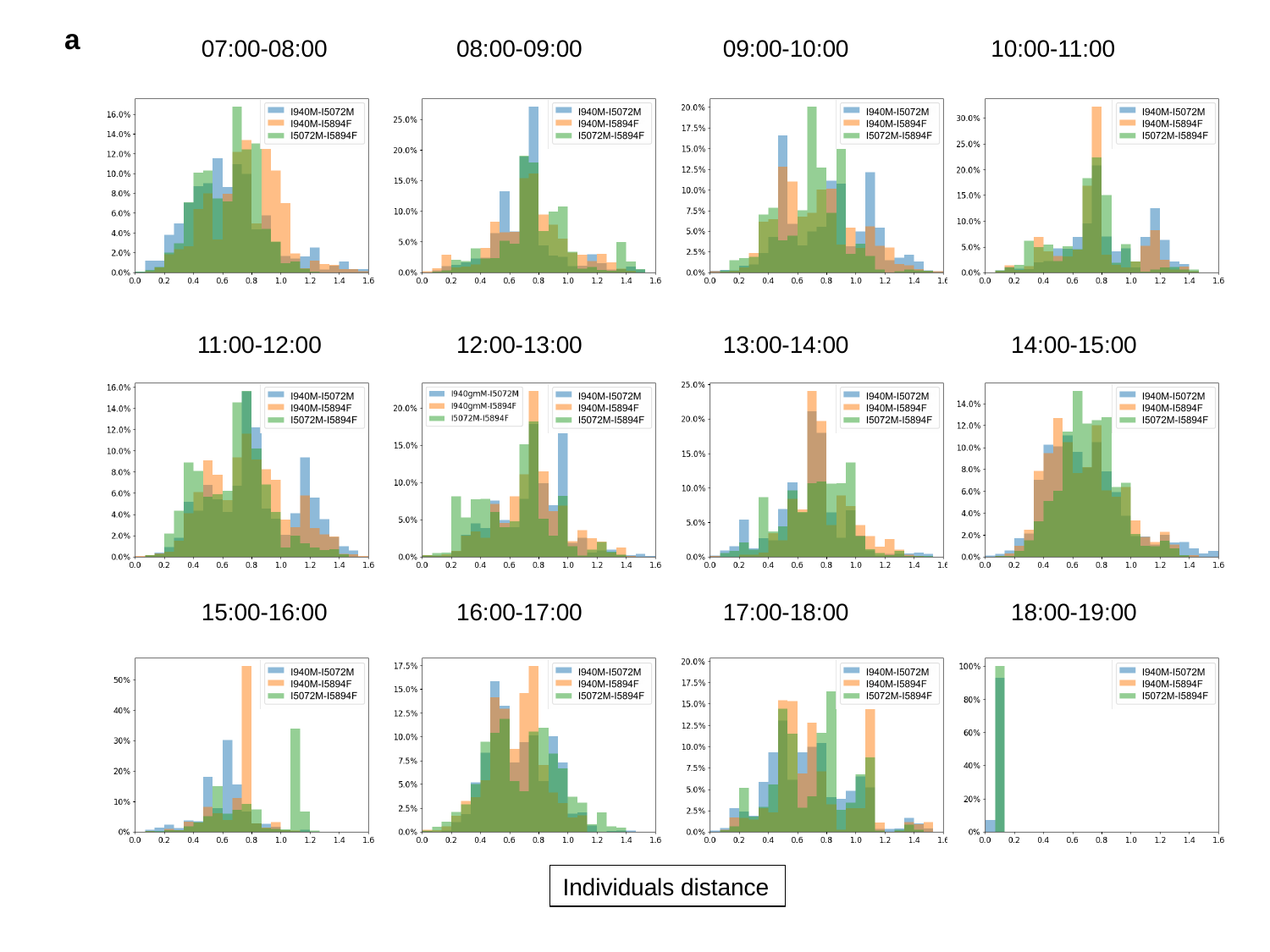

a
07:00-08:00
08:00-09:00
09:00-10:00
10:00-11:00
12:00-13:00
13:00-14:00
14:00-15:00
11:00-12:00
15:00-16:00
16:00-17:00
17:00-18:00
18:00-19:00
Individuals distance

### Slide 10
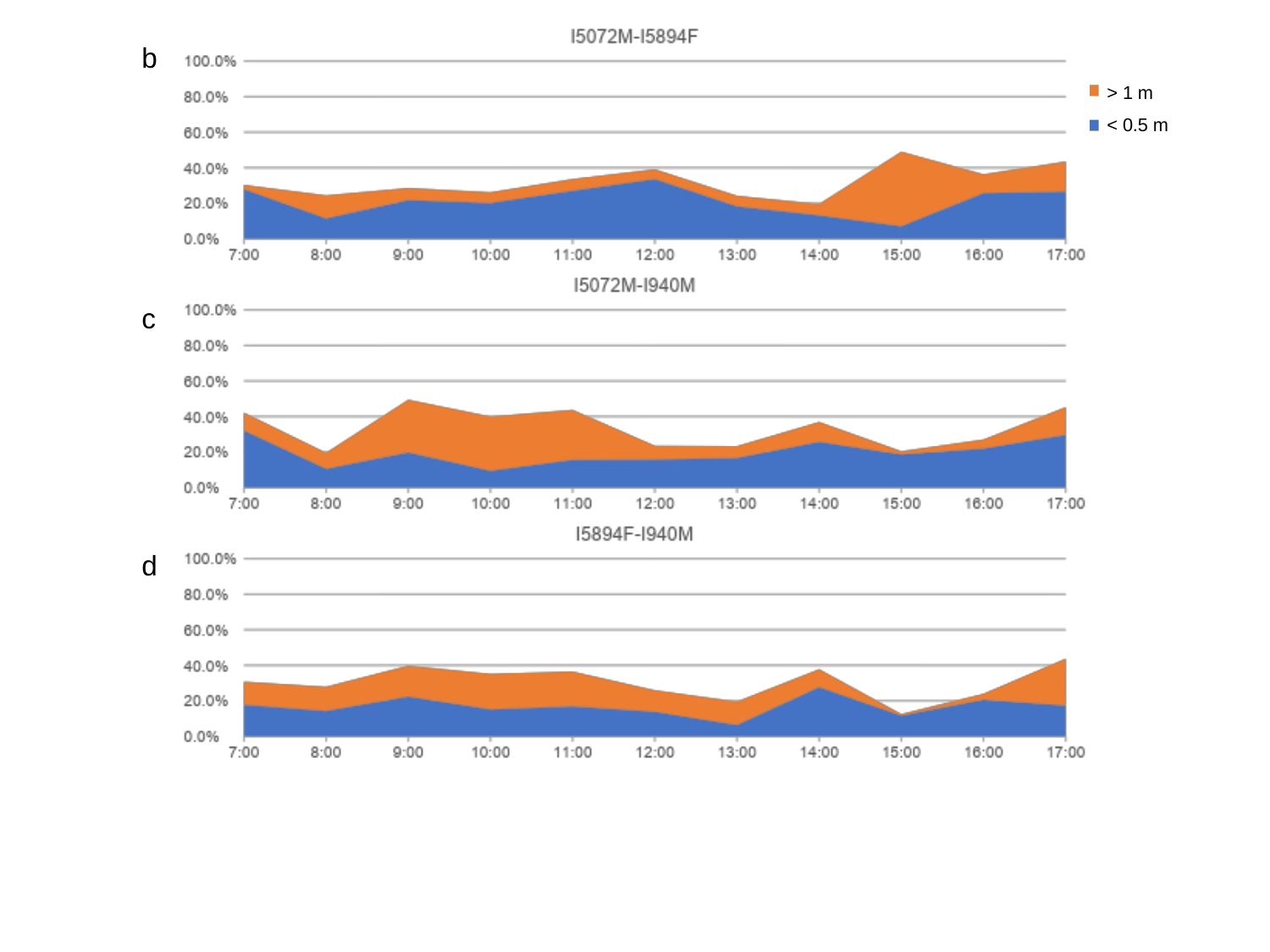

b
> 1 m
< 0.5 m
c
d

### Slide 11
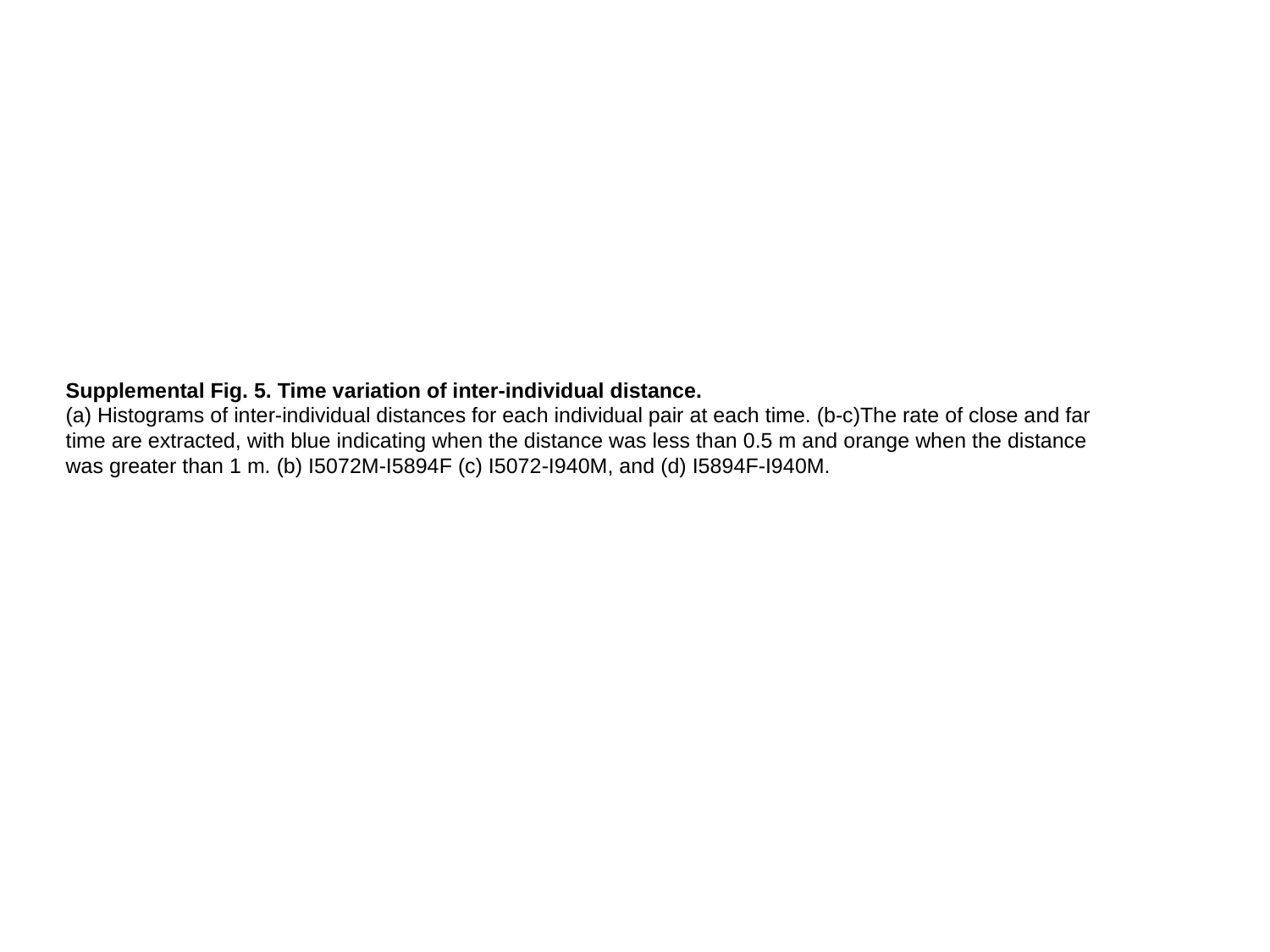

Supplemental Fig. 5. Time variation of inter-individual distance.
(a) Histograms of inter-individual distances for each individual pair at each time. (b-c)The rate of close and far time are extracted, with blue indicating when the distance was less than 0.5 m and orange when the distance was greater than 1 m. (b) I5072M-I5894F (c) I5072-I940M, and (d) I5894F-I940M.

### Slide 12
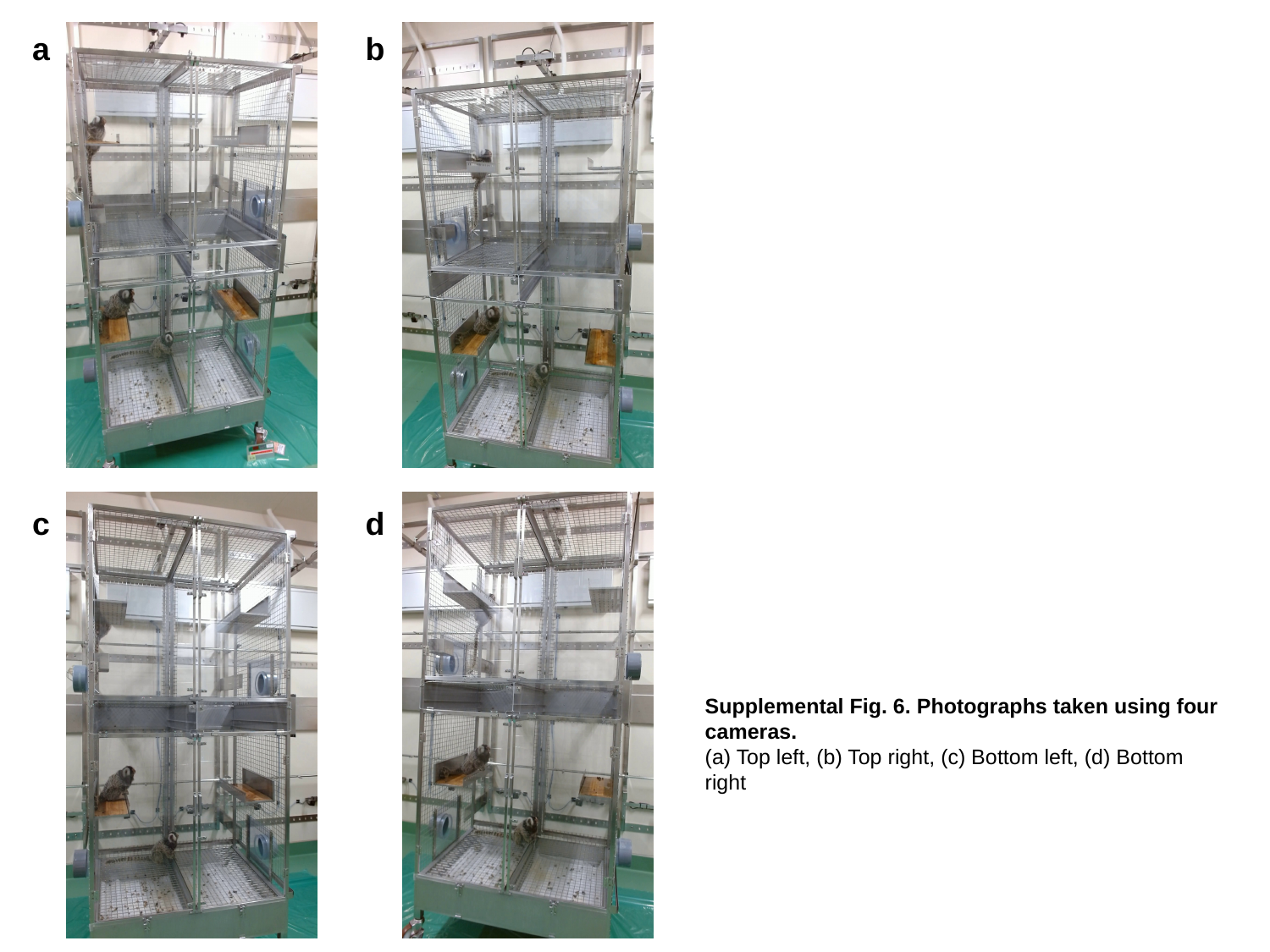

a
b
c
d
Supplemental Fig. 6. Photographs taken using four cameras.
(a) Top left, (b) Top right, (c) Bottom left, (d) Bottom right

### Slide 13
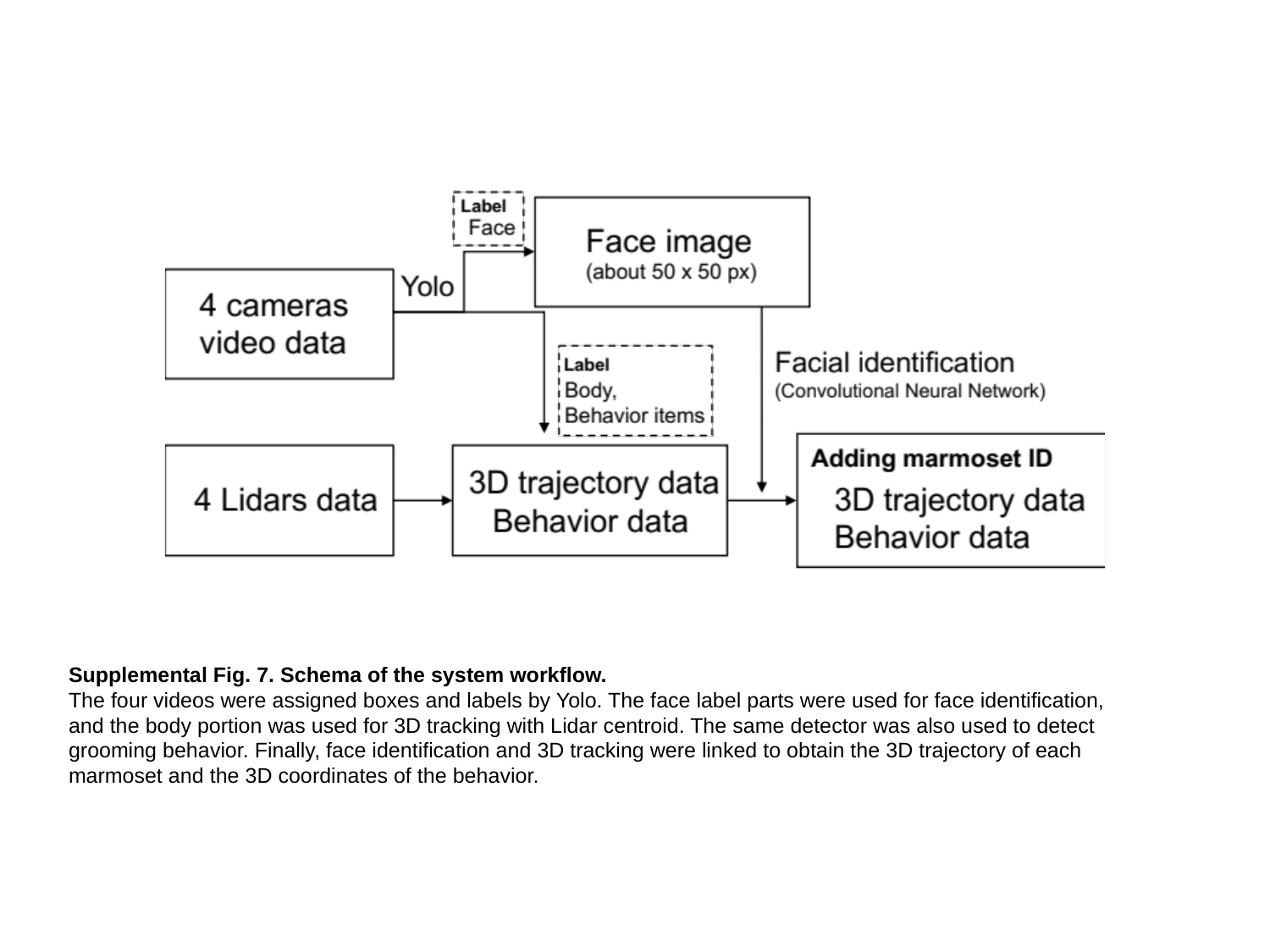

Supplemental Fig. 7. Schema of the system workflow.
The four videos were assigned boxes and labels by Yolo. The face label parts were used for face identification, and the body portion was used for 3D tracking with Lidar centroid. The same detector was also used to detect grooming behavior. Finally, face identification and 3D tracking were linked to obtain the 3D trajectory of each marmoset and the 3D coordinates of the behavior.
