## Supplemental Movie 1 for "Development of a new 3D tracking system for multiple marmosets under free-moving conditions"

### Slide 1
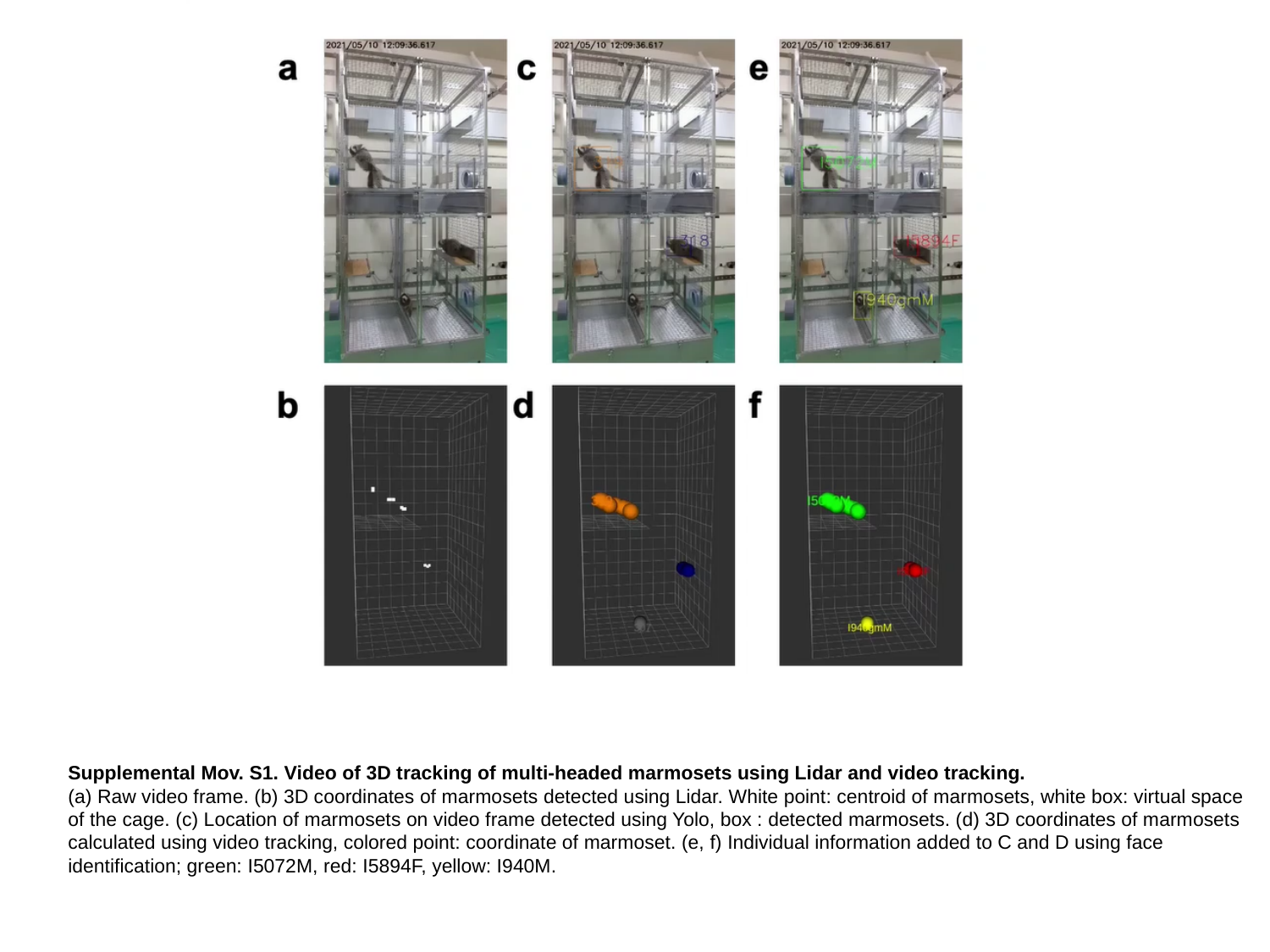

Supplemental Mov. S1. Video of 3D tracking of multi-headed marmosets using Lidar and video tracking.
(a) Raw video frame. (b) 3D coordinates of marmosets detected using Lidar. White point: centroid of marmosets, white box: virtual space of the cage. (c) Location of marmosets on video frame detected using Yolo, box : detected marmosets. (d) 3D coordinates of marmosets calculated using video tracking, colored point: coordinate of marmoset. (e, f) Individual information added to C and D using face identification; green: I5072M, red: I5894F, yellow: I940M.
