## Supplemental Movie 4 for "Development of a new 3D tracking system for multiple marmosets under free-moving conditions"

### Slide 1
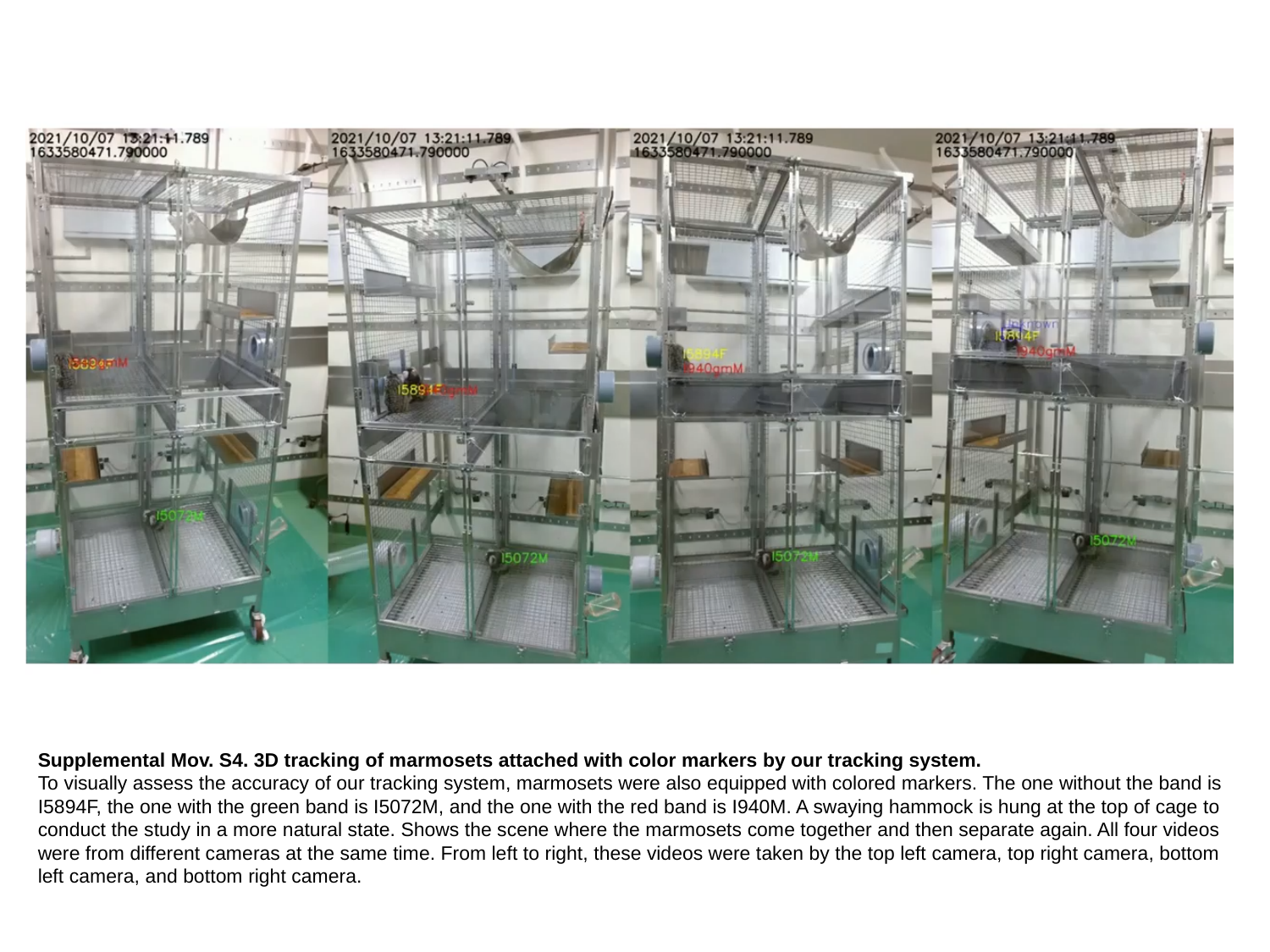

Supplemental Mov. S4. 3D tracking of marmosets attached with color markers by our tracking system.
To visually assess the accuracy of our tracking system, marmosets were also equipped with colored markers. The one without the band is I5894F, the one with the green band is I5072M, and the one with the red band is I940M. A swaying hammock is hung at the top of cage to conduct the study in a more natural state. Shows the scene where the marmosets come together and then separate again. All four videos were from different cameras at the same time. From left to right, these videos were taken by the top left camera, top right camera, bottom left camera, and bottom right camera.
