## Supplemental Movie 5 for "Development of a new 3D tracking system for multiple marmosets under free-moving conditions"

### Slide 1
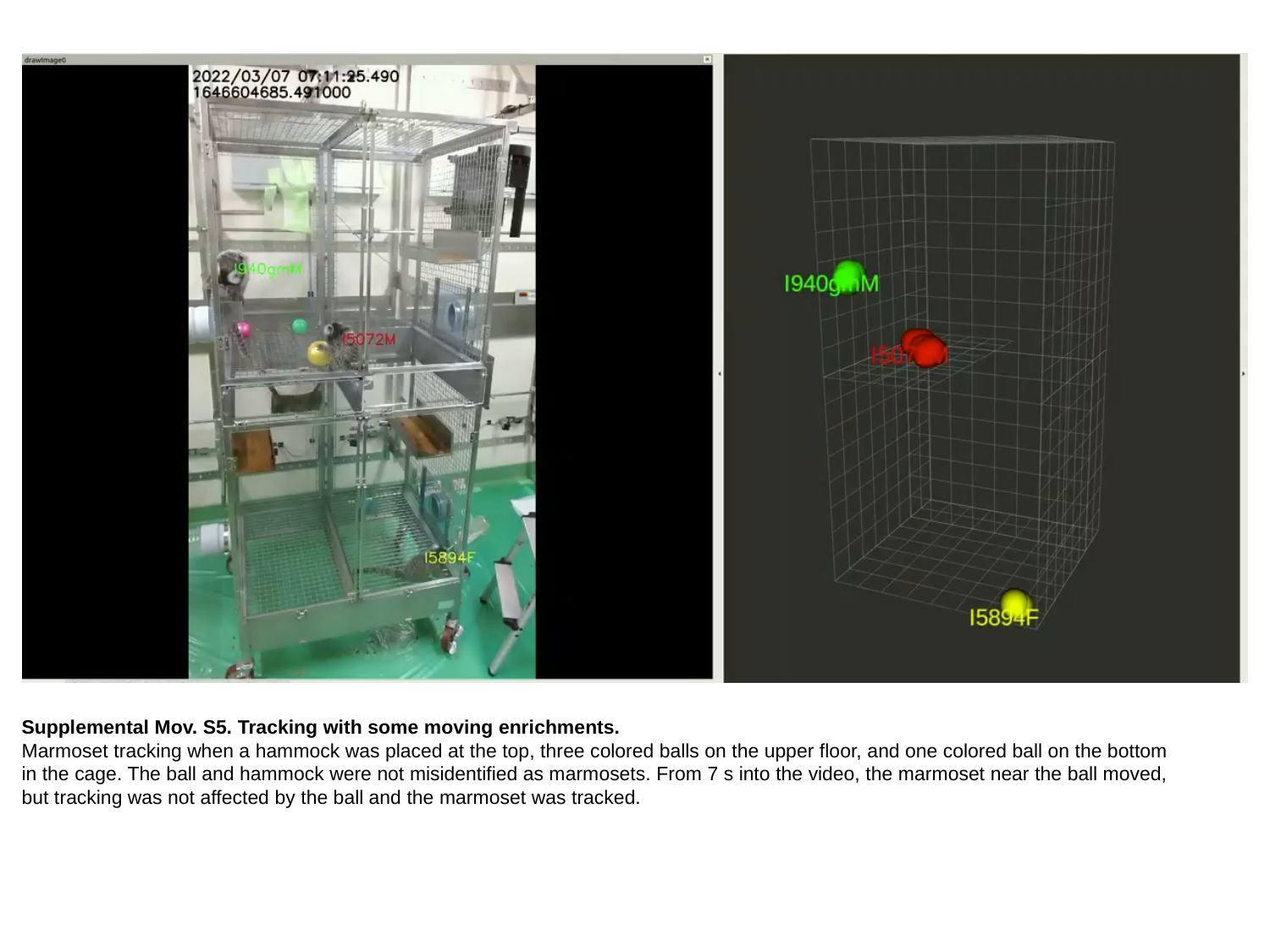

Supplemental Mov. S5. Tracking with some moving enrichments.
Marmoset tracking when a hammock was placed at the top, three colored balls on the upper floor, and one colored ball on the bottom in the cage. The ball and hammock were not misidentified as marmosets. From 7 s into the video, the marmoset near the ball moved, but tracking was not affected by the ball and the marmoset was tracked.
