## Supplemental Movie 6 for "Development of a new 3D tracking system for multiple marmosets under free-moving conditions"

### Slide 1
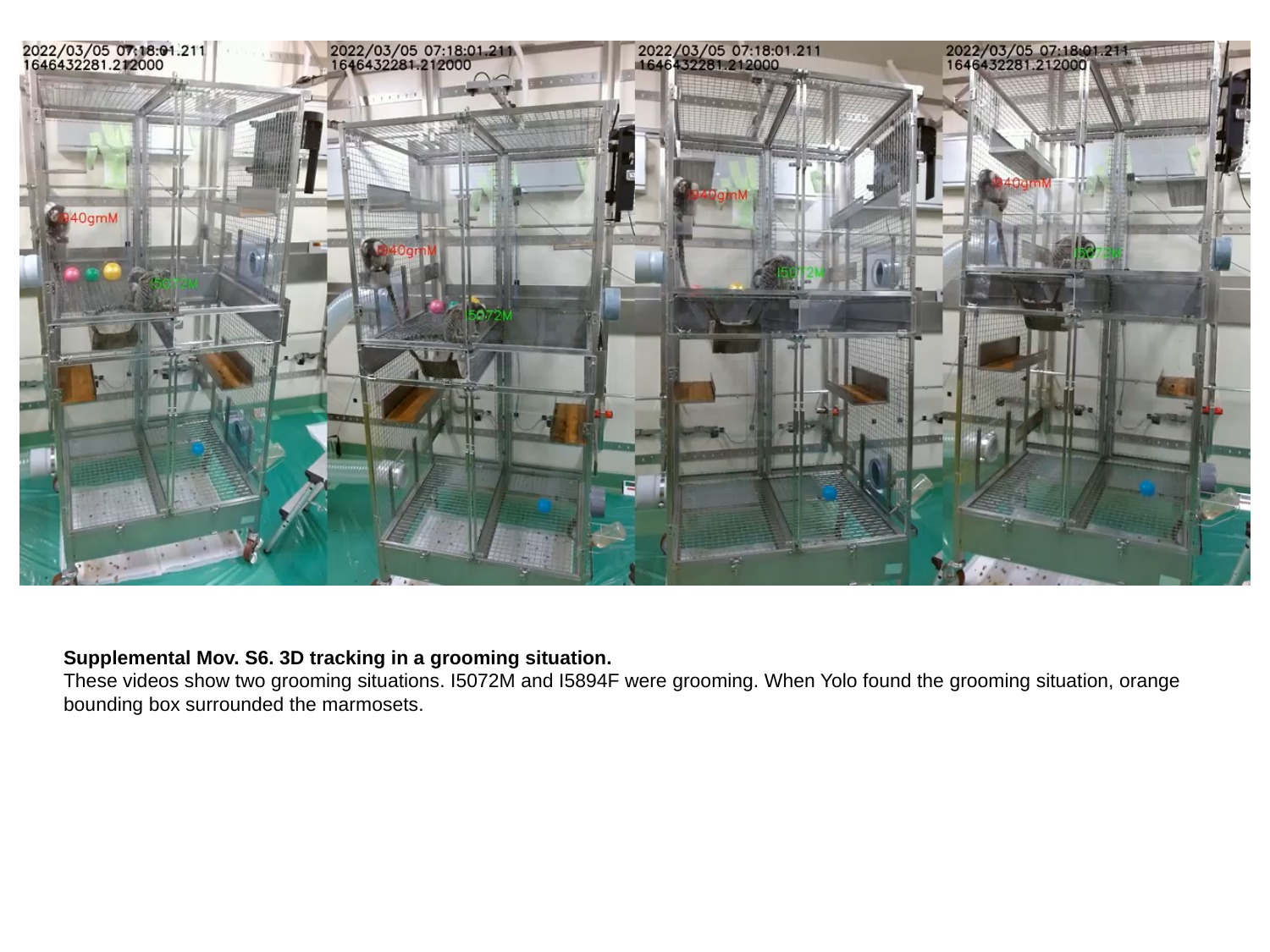

Supplemental Mov. S6. 3D tracking in a grooming situation.
These videos show two grooming situations. I5072M and I5894F were grooming. When Yolo found the grooming situation, orange bounding box surrounded the marmosets.
