## Supplemental Movie 7 for "Development of a new 3D tracking system for multiple marmosets under free-moving conditions"

### Slide 1
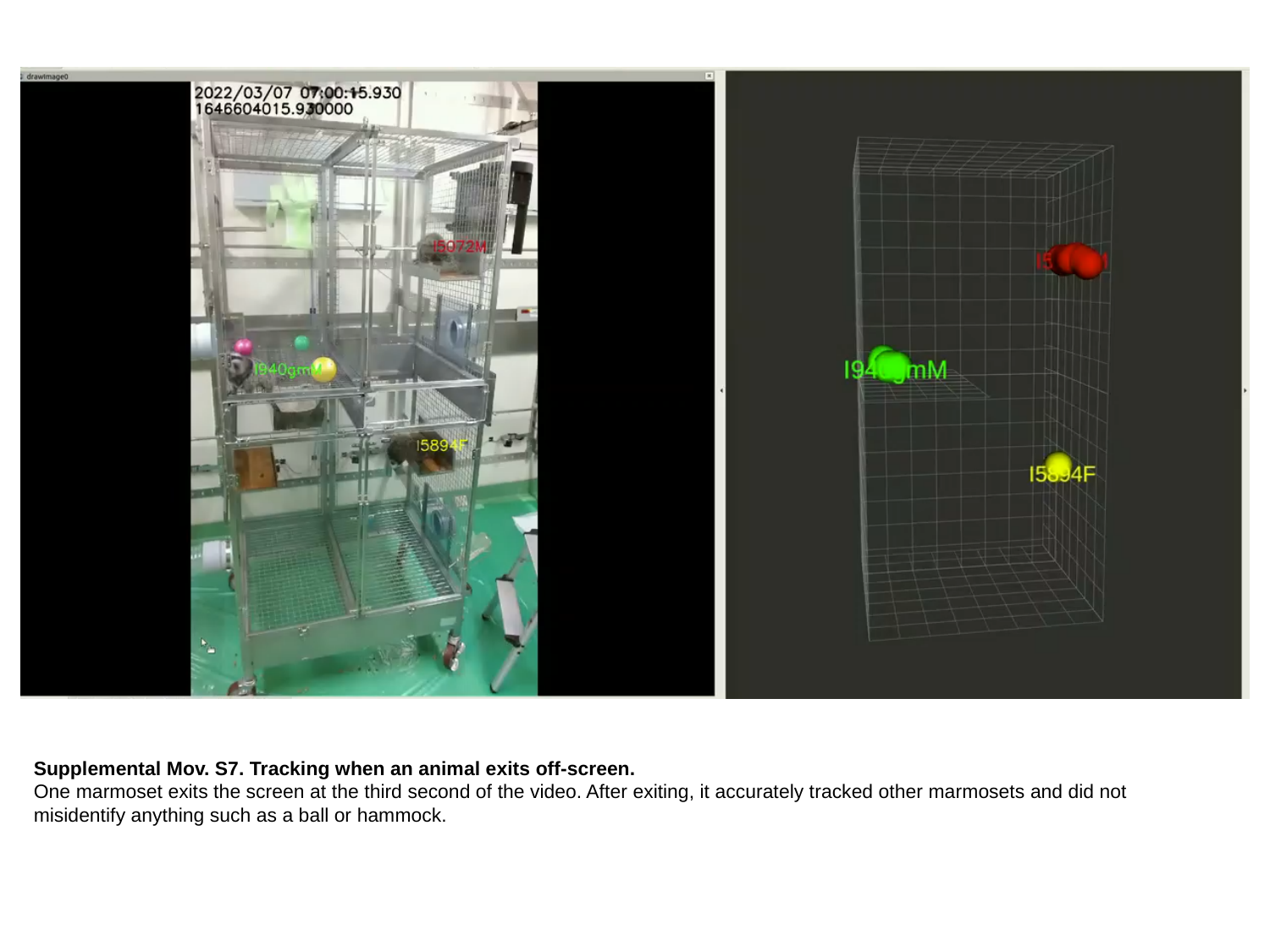

Supplemental Mov. S7. Tracking when an animal exits off-screen.
One marmoset exits the screen at the third second of the video. After exiting, it accurately tracked other marmosets and did not misidentify anything such as a ball or hammock.
