## Supplemental Movie 8 for "Development of a new 3D tracking system for multiple marmosets under free-moving conditions"

### Slide 1
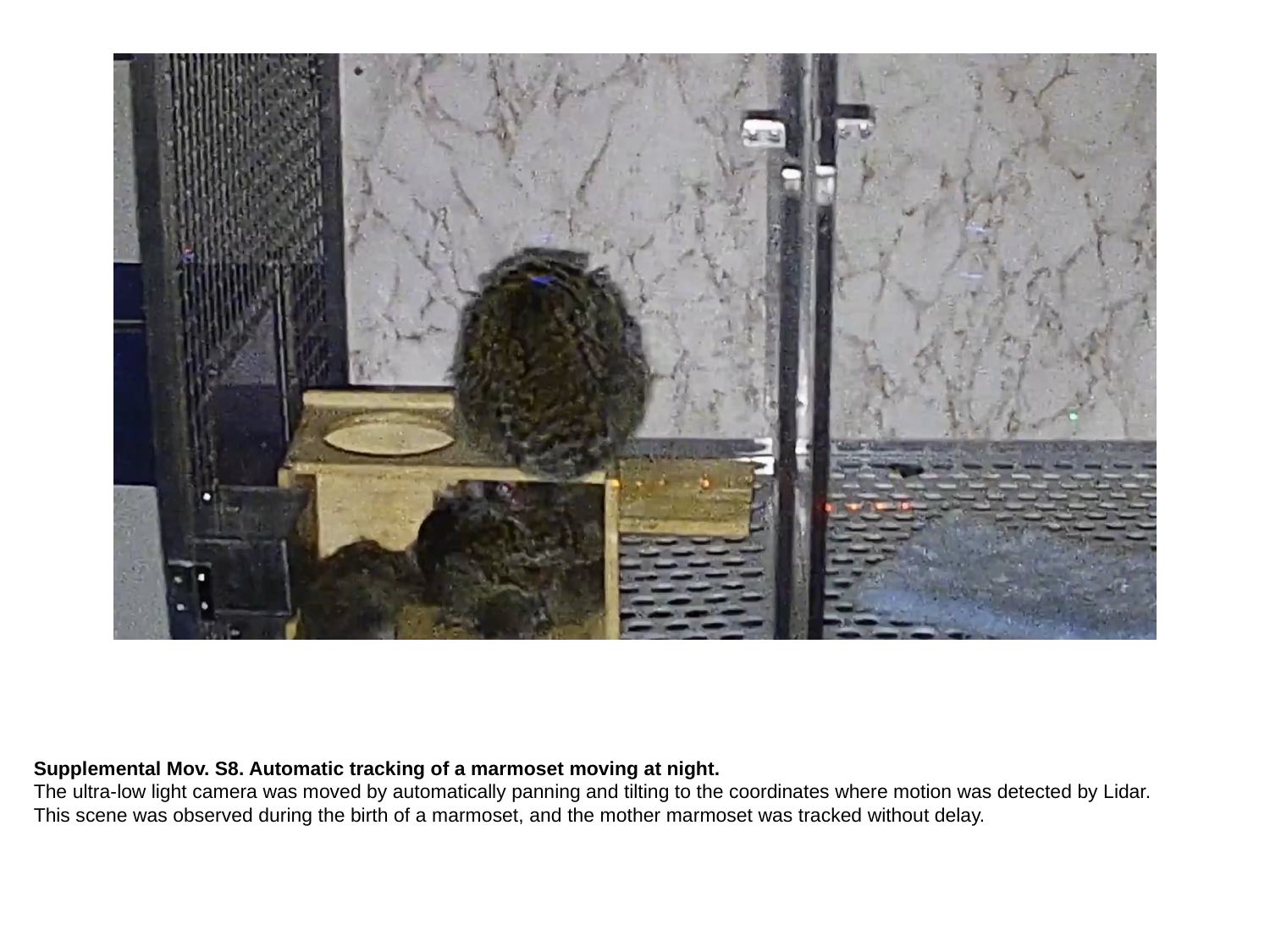

Supplemental Mov. S8. Automatic tracking of a marmoset moving at night.
The ultra-low light camera was moved by automatically panning and tilting to the coordinates where motion was detected by Lidar. This scene was observed during the birth of a marmoset, and the mother marmoset was tracked without delay.
