## Supplemental data 1 for "Development of a new 3D tracking system for multiple marmosets under free-moving conditions"

import datetime

from tensorflow import keras

import numpy as np

import os

import sys

from tensorflow.keras import optimizers

from tensorflow.keras.callbacks import EarlyStopping

import glob

import pandas as pd

import tensorflow.keras

import tensorflow.compat.v1 as tf

from PIL import Image as PILImage

import cv2

import csv

from sklearn.metrics import confusion_matrix

os.environ["TF_FORCE_GPU_ALLOW_GROWTH"]= "true"

def train_face(setting_layer, setting_rate):

print('****train_face****')

#Path Settings

\# train_folder_path = '/home/behavior/Desktop/python/LearnFace/face_image/202109/learning'

write_txt_path = os.path.join(result_folder_path, 'output_class_name_'+ d_today.strftime('%m%d%H%M')+'.txt')

weight_save_path = os.path.join(result_folder_path, 'Marmoset_face_recognition_'+d_today.strftime('%m%d%H%M')+'_vgg19.hdf5')

CLASS_NAME = os.listdir(train_folder_path)

CLASS_NAME.sort()

with open(write_txt_path, 'w') as f_txt:

\# f_txt.write(', '.join(CLASS_NAME))

f_txt.write(str(CLASS_NAME))

#Counting the number of training data and obtaining methods for padding training data

data = str(len(glob.glob(train_folder_path+'/*/*.png')))

image_padding = train_folder_path.split('/')[-1]

#Model settings

model_name = 'vgg19'

IMAGE_SIZE = 224

BATCH_SIZE = 32

fixed_layer = setting_layer

epoch = 50

learning_rate = setting_rate

momentum = 0.9

os.environ["CUDA_VISIBLE_DEVICES"] = "0"

from tensorflow.python.client import device_lib

device_lib.list_local_devices()

\# Read VGG19 model without after FC layer

base_model = keras.applications.vgg19.VGG19(weights='imagenet', include_top=False, input_shape=(224, 224,3))

x = base_model.output

x = keras.layers.GlobalAveragePooling2D()(x)

x = keras.layers.Dense(1024, activation='relu')(x)

predictions = keras.layers.Dense(len(CLASS_NAME), activation='softmax')(x)

model = keras.models.Model(inputs=base_model.input, outputs=predictions)

\# Fixed layer

for layer_no in range(fixed_layer):

layer = model.layers[layer_no]

layer.trainable = False

\# Optimizer

optimizer = keras.optimizers.SGD(lr=learning_rate, momentum=momentum)

\# optimizer = keras.optimizers.Adagrad(lr=learning_rate, epsilon=None, decay=0.0)

\# optimizer = keras.optimizers.Adam(lr=learning_rate, beta_1=0.9, beta_2=0.999, epsilon=1e-08)

model.compile(optimizer=optimizer,

loss='categorical_crossentropy',

metrics=['accuracy'])

model.summary()

train_datagen = keras.preprocessing.image.ImageDataGenerator(

rescale = 1.0 / 255.0,

validation_split=0.2)

test_datagen = keras.preprocessing.image.ImageDataGenerator(

rescale = 1.0 / 255.0,

)

train_generator = train_datagen.flow_from_directory(

train_folder_path,

target_size = (IMAGE_SIZE, IMAGE_SIZE),

batch_size = BATCH_SIZE,

classes = CLASS_NAME,

seed = 1121,

class_mode = 'categorical',

shuffle = True

)

validation_generator = train_datagen.flow_from_directory(

train_folder_path,

target_size = (IMAGE_SIZE, IMAGE_SIZE),

batch_size = BATCH_SIZE,

classes = CLASS_NAME,

seed = 1121,

class_mode = 'categorical',

shuffle = False

)

\# EaelyStopping setting

early_stopping = EarlyStopping(

monitor='val_loss',

\# min_delta=0,

min_delta=1e-5,

patience=2,

mode='auto')

hist = model.fit_generator(

train_generator,

epochs = epoch,

verbose = 1,

validation_data = validation_generator,

workers = 4,

callbacks=[early_stopping]

)

model.save(weight_save_path)

def model_config():

dict = {'model':model_name, 'weight_number':d_today.strftime('%m%d%H%M'), 'learning_rate':str(learning_rate),

'epoch':str(epoch), 'number_of_data':data, 'image_padding':image_padding, 'fixed_layer':str(fixed_layer)}

df = pd.DataFrame(dict,index=['0'])

df.to_csv(result_folder_path+'/model_config.csv')

model_config()

def test_face():

CLASS_NAME = os.listdir(train_folder_path)

CLASS_NAME.sort()

\# setting

IMAGE_SIZE = 224

#CLASS_NAME = ['Unknown', 'I5072M', 'I5894F', 'I940gmM']

os.environ["CUDA_VISIBLE_DEVICES"] = "0"

from tensorflow.python.client import device_lib

device_lib.list_local_devices()

\# Read VGG19 model without after FC layer

base_model = tensorflow.keras.applications.vgg19.VGG19(weights='imagenet', include_top=False)

x = base_model.output

x = tensorflow.keras.layers.GlobalAveragePooling2D()(x)

x = tensorflow.keras.layers.Dense(1024, activation='relu')(x)

predictions = tensorflow.keras.layers.Dense(len(CLASS_NAME), activation='softmax')(x)

model = tensorflow.keras.models.Model(inputs=base_model.input, outputs=predictions)

model.load_weights(weight_path)

graph = tf.get_default_graph()

print('****test_face****')

name_list = ['I5072M', 'I5894F', 'I940gmM','Unknown']

for i in name_list:

id_number = i

\# csv setting

csv_file_name = id_number + 'face_recognition_result' + weight_number+ '.csv'

csv_file_path = os.path.join(result_folder_path, csv_file_name)

csv_f = open(csv_file_path, 'w')

csv_writer = csv.writer(csv_f)

def cv2pil(image):

''' OpenCV -> PIL '''

new_image = image.copy()

new_image = cv2.cvtColor(new_image, cv2.COLOR_BGR2RGB)

new_image = PILImage.fromarray(new_image)

return new_image

image_dir_path = test_img_folder_path + '/' + id_number

globpath = os.path.join(image_dir_path, "*")

image_folder_entry_list_path = glob.glob(globpath) #[a,b,c]

image_folder_entry_list_path.sort()

for image_path in image_folder_entry_list_path:

face_cv2_image = cv2.imread(image_path)

face_image = cv2pil(face_cv2_image)

face_image = face_image.resize((IMAGE_SIZE, IMAGE_SIZE))

face_tensor = tensorflow.keras.preprocessing.image.array_to_img(face_image)

face_tensor = np.expand_dims(face_tensor, axis=0)

face_tensor = tensorflow.keras.applications.vgg19.preprocess_input(face_tensor)

face_tensor = face_tensor/255

\# get face id

preds = model.predict(face_tensor)

probability = preds[0][np.argmax(preds[0])]

probability = '{:.3f}'.format(probability)

#Classification Results

classification_result = CLASS_NAME[np.argmax(preds[0])]

frame_name = image_path.split('/')[-1].split('.')[0]

image_file_name = frame_name + '_' + classification_result + '_' + str(probability) + '.png'

image_write_path = os.path.join(result_folder_path, image_file_name)

#csv

csv_writer.writerow([frame_name, CLASS_NAME[np.argmax(preds[0])], preds[0][np.argmax(preds[0])], id_number])

def result_csv():

print('****result_csv****')

csv_path = result_folder_path + "/accurancy_sum1.csv"

df = pd.DataFrame(columns = [])

for i in glob.glob(result_folder_path + "/*face_recognition_result*"):

tmp = pd.read_csv(i, header=None)

df = pd.concat([df, tmp])

pred_list = df.iloc[:, 1].tolist()

true_list = df.iloc[:, 3].tolist()

cm = confusion_matrix(true_list, pred_list, labels=['I5894F', 'I940gmM', 'I5072M', 'Unknown'])

df = pd.DataFrame(cm)

def confusion_matrix_csv():

pd.DataFrame(cm).to_csv(csv_path,

index = False,

header = ['I5894F', 'I940gmM', 'I5072M', 'Unknown'])

def syuukei():

#ACC1

total = int(df.iat[0,0] + df.iat[0,1] + df.iat[0,2] + df.iat[0,3])

ACC1_I5894F = int(df.iat[0,0])/(total - int(df.iat[0,3]))

ACC1_I940gmM = int(df.iat[1,1])/(total - int(df.iat[1,3]))

ACC1_I5072M = int(df.iat[2,2])/(total - int(df.iat[2,3]))

ACC1_average = (ACC1_I5894F + ACC1_I5072M + ACC1_I940gmM)/3

#ACC2

ACC2_I5894F = int(df.iat[0,0])/total

ACC2_I940gmM = int(df.iat[1,1])/total

ACC2_I5072M = int(df.iat[2,2])/total

ACC2_Unknown = int(df.iat[3,3])/total

ACC2_average = (ACC2_I5894F + ACC2_I940gmM + ACC2_I5072M+ACC2_Unknown) / 4

df_model = pd.read_csv(result_folder_path + "/model_config.csv")

df_model2 = df_model.drop(df_model.columns[0], axis=1)

df_syuukei = df_model2.assign(ACC1_I5894F = ACC1_I5894F, ACC1_I940gmM = ACC1_I940gmM, ACC1_I5072M = ACC1_I5072M, ACC1_average = ACC1_average,

ACC2_I5894F = ACC2_I5894F, ACC2_I940gmM = ACC2_I940gmM, ACC2_I5072M = ACC2_I5072M, ACC2_Unknown=ACC2_Unknown, ACC2_average = ACC2_average)

df_syuukei.to_csv(result_folder_path + "/result.csv", header = True, index = False)

df_syuukei.to_csv('/home/behavior/Desktop/python/LearnFace/4_FaceTest/total_result_20211203.csv', mode='a', header=False, index = False)

confusion_matrix_csv()

syuukei()

#main

layer_list = [9, 12]

learning_rate_list = [1e-3]

train_data_list = ['r',

\# 'rhs', 'rz',

'rzhs']

for g in train_data_list:

train_folder_path = "/home/behavior/Desktop/python/LearnFace/3_TrainFaceImage/6to8/" + g

for i in layer_list:

for j in learning_rate_list:

d_today = datetime.datetime.now()

weight_number = d_today.strftime('%m%d%H%M')

#Path setting

result_folder_path = '/home/behavior/Desktop/python/LearnFace/4_FaceTest'+ '/Result_' + weight_number

test_img_folder_path = '/home/behavior/Desktop/python/LearnFace/3_TrainFaceImage/6to8/original/test_data'

weight_path = result_folder_path + '/Marmoset_face_recognition_' + weight_number +'_vgg19.hdf5'

#Result directory

os.makedirs(result_folder_path, exist_ok=True)

train_face(i, j)

test_face()

result_csv()

print('****learn_face is done.****')
