## Supplemental data 2 for "Development of a new 3D tracking system for multiple marmosets under free-moving conditions"

/**

* Filter by Gaussian weighted moving average over a fixed time slide and window width for a fixed time.\n

* Gaussian weighted moving average processing is performed for each behavior type, and only the behavior data in the time zone exceeding the threshold value is extracted.

* @brief Gaussian weighted moving average filter function

* @param all_behavior_map Behavior detection result information detected by all cameras

* @param FILTER_THRESHOLD Filter threshold

* @param FILTER_GRID_TIME Data subdivision time for filters (unit: seconds)

* @param FILTER_SLIDE_TIME Time to slide the filter range (unit: seconds)

* @param FILTER_WINDOW_TIME Filter window time width (unit: seconds)

* @param file_start_unixtime Video file start time

* @param file_end_unixtime Video file end time

* @return std::map<double, std::vector<behavior_info_csv_divide_writer::BehaviorInfoCsvDivideWriter::Behavior>> Behavior detection result information detected by all cameras after filtering

*/

std::map<double, std::vector<behavior_info_csv_divide_writer::BehaviorInfoCsvDivideWriter::Behavior>> behavior_info_csv_divide_writer::BehaviorInfoCsvDivideWriter::gaussianWeightedMovingAverage(std::map<double, std::vector<behavior_info_csv_divide_writer::BehaviorInfoCsvDivideWriter::Behavior>> all_behavior_map, double FILTER_THRESHOLD, double FILTER_GRID_TIME, double FILTER_SLIDE_TIME, double FILTER_WINDOW_TIME, double file_start_unixtime, double file_end_unixtime){

std::map<double, std::vector<behavior_info_csv_divide_writer::BehaviorInfoCsvDivideWriter::Behavior>> result_all_behavior_filter_map;

// Divide the number of behavior detections into a time range (grid) at regular intervals for each behavior type.

std::map<std::string, std::vector<behavior_info_csv_divide_writer::BehaviorInfoCsvDivideWriter::Grid>> tmp_grid_per_behavior_map = makeGrid(all_behavior_map, FILTER_GRID_TIME, file_start_unixtime, file_end_unixtime);

// Add the acquired grid data

for(auto& itr : tmp_grid_per_behavior_map){

grid_per_behavior_map[itr.first].insert(grid_per_behavior_map[itr.first].end(), itr.second.begin(), itr.second.end());

}

// Pass the grid for each behavior type through a Gaussian weighted moving average filter

unsigned int residue_grid_size = 0;

for(auto& behavior_itr : grid_per_behavior_map){

unsigned int slide_grid_size = std::round(FILTER_SLIDE_TIME / FILTER_GRID_TIME);

unsigned int window_grid_size = std::round(FILTER_WINDOW_TIME / FILTER_GRID_TIME);

unsigned int window_grid_range = window_grid_size;

unsigned int grid_max = behavior_itr.second.size();

double calc_completed_grid_unixtime = 0.0;

for(unsigned int grid_num = 0; grid_num < behavior_itr.second.size(); grid_num++){

// Slide the filter calculation range for a certain period of time

if(grid_num % slide_grid_size == 0){

window_grid_range = grid_num + window_grid_size;

// If the end time of the next X second slide & XX second window exceeds the end time of one video file, the filtering process is interrupted.

if(grid_max <= window_grid_range) break;

// Calculate Gaussian weighted moving average in window range

bool block_f = false;

double old_grid_unixtime = behavior_itr.second[grid_num].grid_unixtime;

double ysum = 0.0;

for(unsigned int calc_window_grid_num = grid_num; calc_window_grid_num < window_grid_range; calc_window_grid_num++){

double diff_grid_unixtime = behavior_itr.second[calc_window_grid_num].grid_unixtime - old_grid_unixtime;

if((FILTER_WINDOW_TIME < diff_grid_unixtime) && block_f == false){

if(old_residue_grid_size == 0) break;

window_grid_range = old_residue_grid_size - 1;

block_f = true;

}

old_grid_unixtime = behavior_itr.second[calc_window_grid_num].grid_unixtime;

// Gaussian weighted moving average calculation

double x = ((grid_num + FILTER_WINDOW_TIME) - calc_window_grid_num) / FILTER_WINDOW_TIME * 3.0;

double y = std::exp(-1 * std::pow(x, 2) / 2) / std::sqrt(2 * M_PI); // Standard normal distribution

ysum = ysum + y * behavior_itr.second[calc_window_grid_num].behavior_count;

}

unsigned int next_slide_grid_num = grid_num + slide_grid_size;

if(grid_max <= next_slide_grid_num) next_slide_grid_num = grid_max -1;

// Save the behavior when the filter value exceeds the threshold

if(FILTER_THRESHOLD <= ysum){

for(unsigned int get_grid_data_num = grid_num; get_grid_data_num < next_slide_grid_num; get_grid_data_num++){

for(auto& data_itr : behavior_itr.second[get_grid_data_num].behavior_map){

result_all_behavior_filter_map[data_itr.first].insert(result_all_behavior_filter_map[data_itr.first].end(), data_itr.second.begin(), data_itr.second.end());

}

}

}

calc_completed_grid_unixtime = behavior_itr.second[next_slide_grid_num - 1].grid_unixtime;

}

}

// Delete unnecessary grid data

for(int clear_grid_num = behavior_itr.second.size() - 1; 0 <= clear_grid_num; clear_grid_num--){

if( behavior_itr.second[clear_grid_num].grid_unixtime <= calc_completed_grid_unixtime ){

behavior_itr.second.erase(behavior_itr.second.begin() + clear_grid_num);

}

}

residue_grid_size = behavior_itr.second.size();

}

old_residue_grid_size = residue_grid_size;

return result_all_behavior_filter_map;

}
